# Synthesis and Evaluation of a Stable Isostere of Malonyllysine

**DOI:** 10.1101/2020.08.23.263285

**Authors:** Sarah E. Bergholtz, Yihang Jing, Rhushikesh A. Kulkarni, Thomas T. Zengeya, Jordan L. Meier

**Author notes:** These authors contributed equally (Sarah E. Bergholtz, Yihang Jing, Rhushikesh A. Kulkarni).

## Abstract

Lysine malonylation is a recently characterized posttranslational modification involved in the regulation of energy metabolism and gene expression. Two unique features of this posttranslational modification are its negative charge and potential susceptibility to decarboxylation, both of which pose possible challenges to its study. As a step towards addressing these challenges, here we report the synthesis and evaluation of a stable isostere of malonyllysine. First, we find that synthetic substitution of the malonyl group with a tetrazole isostere results in amino acids resistant to thermal decarboxylation. Next, we demonstrate that protected variants of this amino acid are readily incorporated into peptides. Finally, we show that tetrazole isosteres of malonyllysine can be recognized by anti-malonyllysine antibodies, validating their ability to mimic features of the endogenous lysine modification. Overall, this study establishes a new chemical strategy for stably mimicking a metabolite-derived posttranslational modification, providing a foothold for tool development and functional analyses.

## Introduction

Reversible acetylation plays a critical role in many important cellular functions, including the regulation of metabolism and gene expressions.^1^ This paradigm has been further expanded by the identification of several related posttranslational modifications, collectively referred to as lysine acylations.^2–3^ Similar to acetylation, the majority of lysine acylations are derived from acyl-CoA metabolites and have been found to be enzymatically removed. However, many of these modifications are physiochemically distinct from acetylation, allowing them to exert unique effects on the properties of modified proteins. One of the most distinct is lysine malonylation (Fig. 1a). Lysine malonylation is a posttranslational modification first discovered in 2011 via LC-MS/MS analysis of bacterial and cancer cell proteomes.^4^ Malonylation possesses a larger steric footprint than acetylation and also exerts a more profound electrostatic effect, replacing the positively charged epsilon amine of protein lysine residues with a negatively charged carboxylate. Lysine malonylation is derived from malonyl-CoA, a precursor for anabolic fatty acid synthesis whose production is highly regulated by the metabolic state of the cell.^5^ In addition to being intrinsically sensitive to fed/fasted state, levels of malonyl-CoA can be altered by cellular transformation, administration of fatty acid synthase inhibitors, and inborn errors of metabolism known as malonic acidurias.^6^ Removal of lysine malonylation is catalyzed by a SIRT5, an NAD-dependent deacylase enzyme which primarily localizes to the mitochondria but can also be found in the cytosol and nucleus.^4, 7^ In contrast, protein acetyltransferase enzymes that catalyze lysine malonylation at appreciable rates have yet to be identified. Building on early observations of non-enzymatic malonylation, our group recently applied a reactivity-based approach to define malonyl-CoA as a hyperreactive metabolite that drives non-enzymatic lysine malonylation in the cytosol and nucleus.^8^ Non-enzymatic malonylation was found to functionally alter glycolytic enzyme activities, consistent with the biological observations of others.^9^ These findings hint the at potential of lysine malonylation to both influence signaling, via effects on protein structure and activity, as well as significantly reflect an organism’s history of environmental exposures and metabolic state.

**Figure 1.**
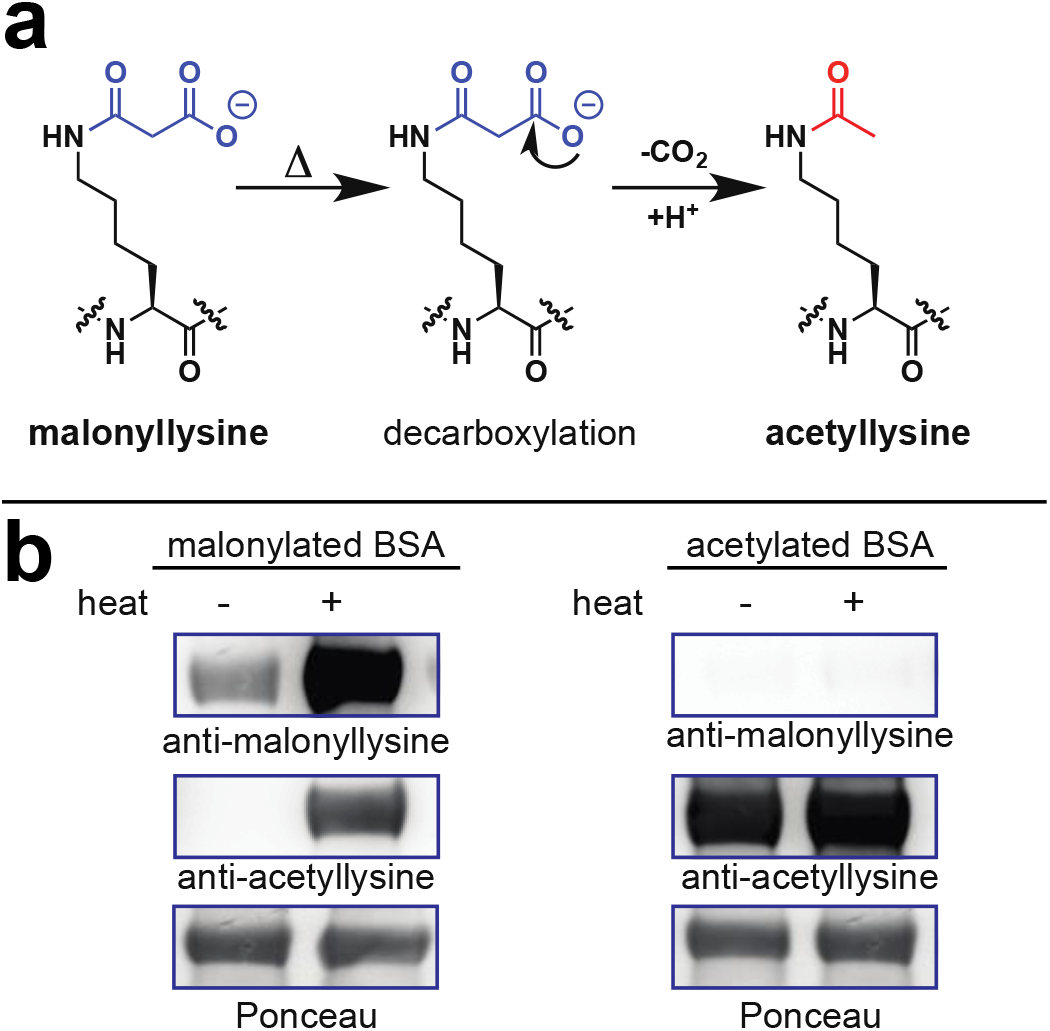
(a) Decarboxylation of malonyllysine to acetyllysine. (b) No decarboxylation of malonyllysine is seen at room temperature. However, heat can cause chemical decarboxylation of malonyllysine-containing proteins to form acetyllysine, a cross-reactive epitope.

One relatively uncharacterized property of lysine malonylation is its potential chemical lability. Upon heating malonyl amides are known to undergo decarboxylation to acetamides in solution, which has the effect of converting malonyllysine to acetyllysine within proteins (Fig. 1a). This reaction was first noted in early studies of protein malonylation, which used the facile decarboxylation of malonyllysine-containing peptides in the gas-phase as a characteristic signature for identification of malonyllysine in multi-stage peptide mass spectrometry experiments.^4^ This reactivity is unique to lysine malonylation versus other acylations, and is conceptually similar to labile phosphorylations such as phosphohistidine, phosphoarginine, and phospholysine, whose instability has been found to hinder their detection and investigation.^10^ In contrast, the ability to detect and raise antibodies against malonllysine suggests its reactivity may be relatively modest compared to labile phosphorylations, and may only become relevant under specific conditions.

In order to better characterize the potenttial chemical lability of malonyllysine, we first set out to generate a malonylated protein standard for use in model reactions. In previous studies, we have applied acyl-CoAs as well as N-acetyl-cysteamine (NAC) esters as non-enzymatic acylation reagents.^8^ To prepare malonylated bovine serum albumin (BSA) for use in model studies, we thus compared three non-enzymatic malonylation agents: malonyl-CoA, malonyl-NAC, and Meldrum’s acid, which can malonylate proteins via a decarboxylative acylation reaction (Fig. S1a). Reagents were individually incubated with BSA overnight, and analyzed for malonylation via SDS-PAGE and anti-malonyllysine immunoblotting. The relative intensity of malonylation in these treated samples was found to follow the order: malonyl-NAC > malonyl-CoA >> Meldrum’s acid (Fig. S1b). These studies indicate malonyl-NAC as an effective reagent for in vitro non-enzymatic protein malonylation. Next, we explored our samples for evidence of decarboxylation. In contrast to its sensitive detected by anti-malonyllysine Western blot, malonylated BSA possessed no observable cross-reactivity with acetyllysine antibodies (Fig. 1b), indicating it is relatively stable at room temperature. However, heating of malonylated BSA under conditions commonly employed for SDS-PAGE loading (90 °C, 5 min) resulted in production of an intense acetylation signal (Fig. 1b). These studies suggest malonyllysine is a stable posttranslational modification under physiological conditions which demonstrates “cryptic” chemical lability upon heating.

Studies of labile posttranslational modifications such as phosphohistidine have greatly benefited from the development of stable analogues of these modifications.^10^ In this approach, synthetic amino acids are developed that mimic the chemical properties of the native posttranslational modification but which possess sufficient stability to allow for incorporation into proteins and peptides for functional analysis. Access to such amino acids can facilitate antibody generation as well as the interrogation of receptor-binding interactions dependent upon the modified residue, enabling fine-tuning of affinity and selectivity. This notion inspired us to ask the question: could one design an isostere of malonyllysine? As an initial proof-of-concept, we explored the effect of replacing the terminal carboxy-group of malonyllysine with 5-substituted-1H-tetrazoles.^11^ 5-substituted tetrazoles (also referred to herein as tetrazolic acids) are a class of heterocyclic bioisosteres that exhibit similar physical properties to carboxylic acids. These include a planar structure as well as a nearly identical pKa, which leads to similar ionization at physiological pH. Distinct chemical features of tetrazoles include the delocalization of negative charge around the ring, as well as relatively increased size and lipophilicity (Fig. 2a). Most importantly, unlike the terminal carboxy-group found in malonyllysine, tetrazoles cannot decompose via entropically favorable release of carbon dioxide. These considerations led us to evaluate 5-substituted tetrazoles as a stable class of malonyllysine isosteres.

**Figure 2.**
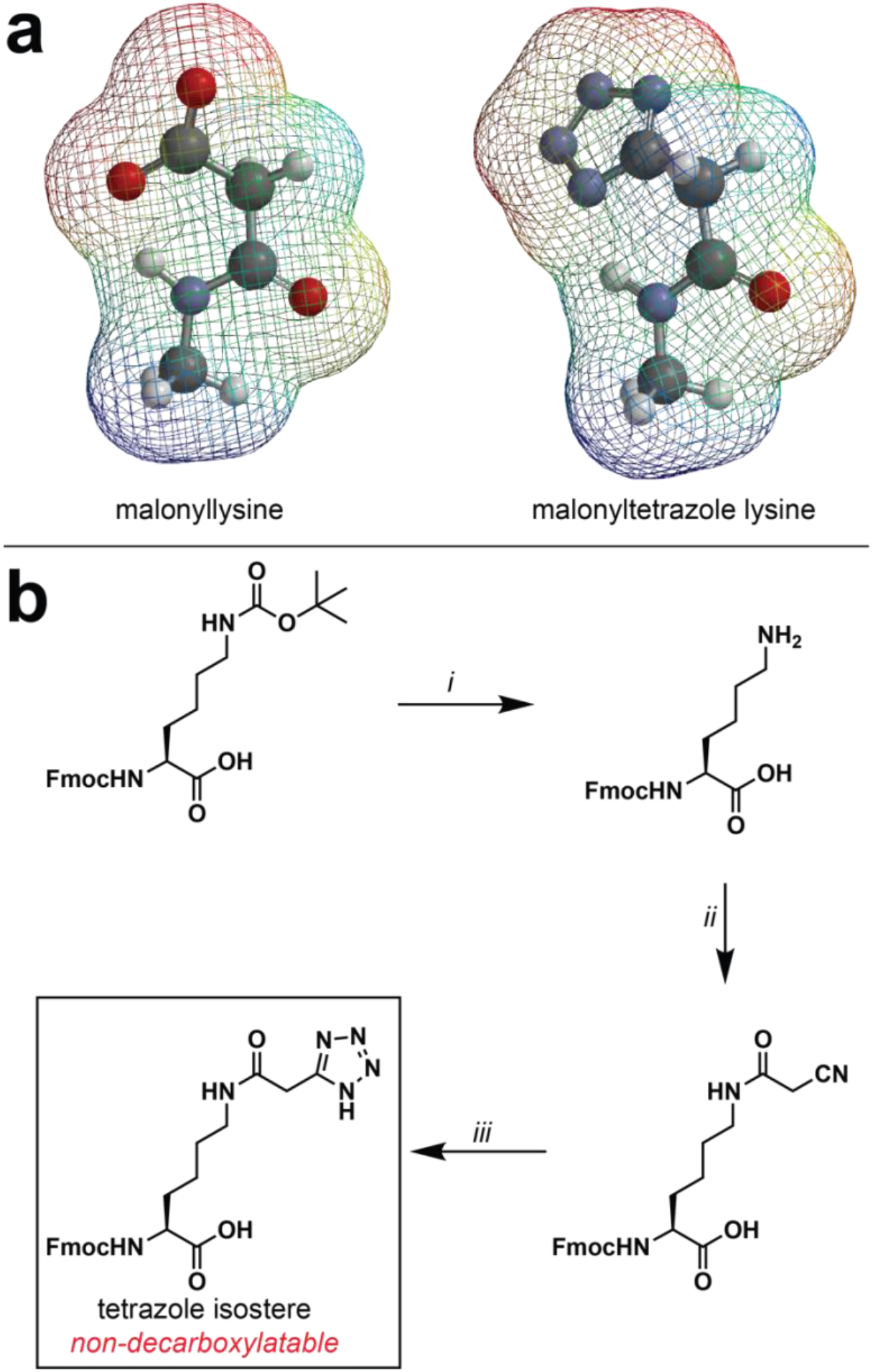
(a) Electrostatic potential map comparing malonyllysine and a malonyltetrazole lysine isostere. (b) Synthesis of a tetrazole isostere of malonyllysine amino acid.

Tetrazole mimics of malonyllysine were prepared via a facile three-step synthesis from the commercially available Fmoc-protected lysine amino acid (Fig. 2b). Following deprotection, cyanoacetic acid was coupled to the epsilon amine of Fmoc-Lys-OH using standard carbodiimide-mediated coupling conditions. Pilot studies found that omitting bases such as diisopropylethylamine from this reaction led to greatly improved yields, possibly by limiting decomposition of the highly acidic O-acylisourea of cyanoacetic acid. An alternative route, in which lysine cyanoacetamide was produced via sequential bromoacetic acid coupling and cyanide displacement, was also explored and found to provide product, albeit with reduced yields. The lysine cyanoacetamide was further subjected to zinc bromide-catalyzed cycloaddition with sodium azide to furnish the 5-substituted tetrazole amino acid. These studies establish a straightforward synthetic route to tetrazole isosteres of a malonyllysine.

In drug design, useful isosteres mimic the parent functional group while simultaneously imbuing a biomolecule with novel properties, such as increased binding affinity or metabolic stability. To establish the distinct chemical properties of tetrazole and malonyllysine amino acids, we compared their susceptibility to the aforementioned thermally-induced decarboxylatve decomposition. For these studies, Fmoc-protected malonyllysine and malonyltetrazole amino acids were incubated in neutral PBS (pH 7) for 0-12 hours at temperatures ranging from 23 °C to 90 °C. As anticipated from our studies of malonylated BSA, incubation of the malonyllysine amino acid at elevated temperatures resulted in increased decarboxylation, as assessed by the relative ion intensity of malonylated versus to acetylated amino acids (Fig. 3). In contrast, no similar decomposition was observed with the malonyltetrazole amino acid. These observations indicate 5-substituted malonyltetrazoles may serve as chemically stable alternatives to cryptically labile malonylated amino acids.

**Figure 3.**
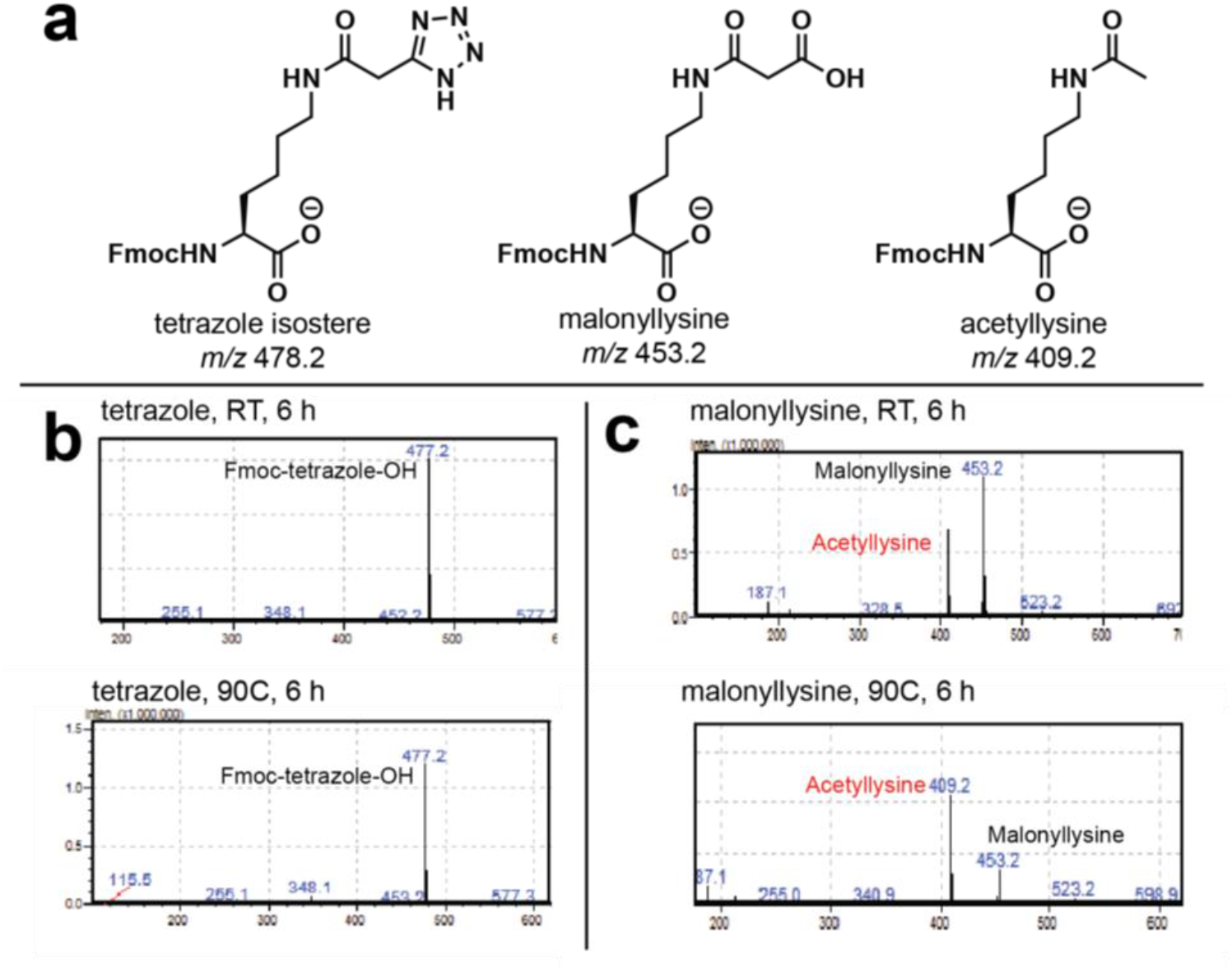
(a) Calculated masses of modified lysine amino acids. (b) Fmoc-malonyltetrazole lysine is stable to prolonged heating at 90 °C. (c) Fmoc-malonyllysine decarboxylates upon ionization, and increasing levels of the decarboxylated mass are observed upon analysis of amino acids which have been heated at 90 °C.

Next, we explored whether malonyltetrazole amino acids could be incorporated into biologically-relevant sites of malonylation using conventional solid-phase peptide synthesis. As a target for these studies, we chose a peptide based on the N-terminus of the nuclear structural protein histone H2B (Fig. 4a). Recent proteomic studies found that malonylation of lysine 5 in histone H2B undergoes a large change in abundance upon knockout of SIRT5, suggesting it is highly regulated.^9^ Coupled with the known role of histone acylation in chromatin structure and transcription,^1^ this suggests H2K5 may be an important node linking malonylation with gene expression in the nucleus. Considering building blocks for the introduction of malonyl isostere, we opted to utilize **1** without any further protection of the acidic tetrazole moiety. We reasoned the lack of an additional protection/deprotection step would be preferable from a technical perspective, as it may facilitate the production of sufficient quantities of amino acid to allow its utilization in excess during solid-phase peptide synthesis reactions. Previous studies have found unprotected 5-substituted tetrazoles to be compatible with peptide coupling reagents,12 suggesting the synthetic feasibility of this strategy. Accordingly, peptides based on histone H2B (residues 1-11) containing either malonyltetrazole or native lysine at K5 were synthesized according to standard Fmoc solid-phase peptide synthesis methods on Wang resin (Fig. 4a). Cleavage from resin with TFA followed by reverse-phased HPLC purification provided the cognate peptides in multimilligram amounts. These studies demonstrate the compatibility of malonyltetrazole amino acids with solid-phase synthesis strategies.

**Figure 4.**
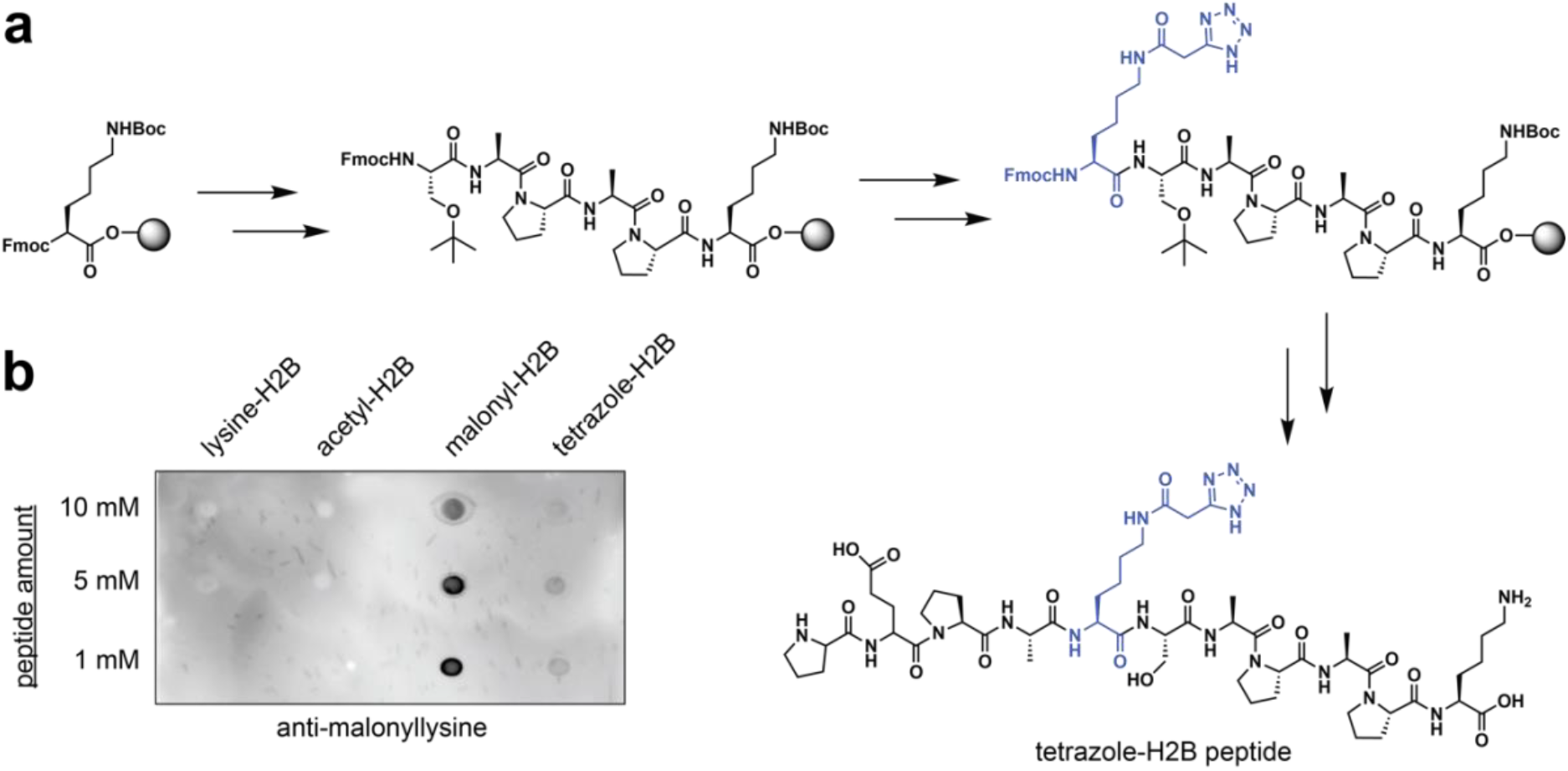
Incorporation of tetrazole isosteres of malonyllysine into peptides by solid-phase peptide synthesis. (a) Synthetic scheme. (b) Peptides containing tetrazole isosteres are recognized by anti-malonyllysine antibodies.

Having established a unique chemical property of malonyltetrazole, as well as the ability to incorporate it into peptides, one final question was whether this non-native amino acid could mimic the interaction of malonyllysine with biomolecules. To assess this, we tested the recognition of these two amino acids using anti-malonyllysine antibodies by dot blot. These studies utilized the aforementioned H2B (1-11) peptides derived at position 5 with either lysine, acetyllysine, malonyllysine, or malonyltetrazole Peptides were spotted onto nitrocellulose membranes, blocked, and probed with a polyclonal anti-malonyllysine antibody (PTM Biolabs #PTM-901) followed by secondary detection.^4^ These dot blot experiments revealed the polyclonal antibody readily detects malonyllysine and malonyltetrazole, but not acetyllysine or unmodified lysine, in the H2B peptide context (Fig. 4b). This suggests malonyltetrazole- and malonyllysine-containing peptides can present similar molecular interaction surfaces for receptor recognition. Interestingly, a significantly less intense signal was observed for the malonyltetrazole H2B peptide compared to the analogous malonyllysine species. This suggests that within the polyclonal anti-malonyllysine antibody population, individual antibodies exist that recognize conformations of malonyllysine not mimicked by the tetrazole analogue. Consistent with this, using dot blot analysis we identified one monoclonal anti-malonyllysine antibody (CST #) that is able to distinguish malonyllysine and malonyltetrazole (Fig. S2).^7^ Distinct chemical features of malonyltetrazole that may preclude detection by this antibody include equilibration to the 2H-5-substituted tetrazole isomer, as well as increased size, lipophilicity, and delocalization of negative charge. Overall, these findings indicate the receptor interaction landscape presented by malonyltetrazole- and malonyllysine-is characterized both by overlapping, as well as distinct, molecular recognition surfaces.

To summarize, here we have reported the design and characterization of a stable bioisostere of malonyllysine. Lysine amino acids incorporating a 5-substituted malonyltetrazole at the epsilon amino group serve as synthetically accessible analogues of malonyllysine. Model studies indicate this analogue possesses reduced thermal lability relative to the native malonyl-posttranslational modification, and can be readily incorporated into peptides via solid-phase peptide synthesis. Finally, we utilized antibody-based dot blotting to demonstrate the ability of malonyltetrazole-containing peptides to present a molecular interaction surface which overlaps with that of malonyllysine. The knowledge generated from this proof-of-concept study should find several applications. First, the cryptic chemical lability of malonyllysine has to date been only sparingly characterized in the literature.^4^ Our results using malonylated BSA indicate that while malonyllysine is quite stable at room temperature, but that studies of this posttranslational modification should take care to avoid heat, a condition proteins are routinely exposed to during preparation for denaturing polyacrylamide gel electrophoresis. This decarboxylation of malonyllysine has the potential not only to limit detection of this malonyllysine, but also to cause artifactual detection of acetyllysine at authentic malonylation sites. Second, our studies suggest that in cases where the terminal carboxy-functional group of malonyllysine hinders applications, malonyltetrazoles may be employed as alternative bioisostere that are easily incorporated into peptides and engage in similar receptor binding interactions as the natural modified amino acid. One caveat to this strategy stems from our identification of a monoclonal antibody that distinguishes malonyllysine and malonyltetrazole, which suggests malonyltetrazole may only partially mimic the molecular interaction landscape presented to the immune system by malonyllysine. Whether the converse is true, i.e. can malonyltetrazole elicit the production of antibodies that do not recognize malonyllysine, remains unknown but will be important to explore if this amino acid is employed as an immunogen. A multitude of carboxylate isosteres are known in the synthetic literature, and an interesting future question is whether replacing the tetrazole with one of these alternatives may more effectively mimic malonyllysine and circumvent this limitation. It is important to note that, as we explicitly note in the text, unlike phospho-amino acids the generation of malonyllysine antibodies is not necessarily limited hindered by their cryptic lability, and further motivation for asking whether effective isosteres of this modification can be generated lies in their potential to enable additional applications. The ability to incorporate malonyltetrazole into peptides, as well as possibly into proteins via native chemical ligation,10 may enable the identification of proteins which interact with this modification with high affinity as ‘readers.’ The increased lipophilicity, cell permeability, and “cage-ability” of tetrazoles relative to carboxylates may also usefully incorporate this analogue into proteins via unnatural amino acid mutagenesis approaches.^13^ Finally, these studies open the door to the development of bioisostere of additional dicarboxylate metabolite-derived lysine acylations, including lysine succinylation and glutarylation.^6^ In particular it will be interesting to evaluate whether (and how) these analogues influence turnover by SIRT-family deacylases. By expanding the methodological arsenal available to study reversible acylation, these new tools may facilitate a greater understanding of how posttranslational modifications function as key mediators of biology and disease.

## Supporting Information

Supporting Information including detailed materials and methods, as well as supplemental figures is available online.

## Acknowledgements

We thank Arissa Bavari for contributing synthetic procedures for this study. This work was supported by National Institutes of Health, National Cancer Institute, Center for Cancer Research (ZIABC011488).

## References

1. Ali, I.; Conrad, R. J.; Verdin, E.; Ott, M., Lysine Acetylation Goes Global: From Epigenetics to Metabolism and Therapeutics. Chem Rev 2018, 118 (3), 1216–1252.

2. Lin, H.; Su, X.; He, B., Protein lysine acylation and cysteine succination by intermediates of energy metabolism. ACS Chem Biol 2012, 7 (6), 947–60.

3. Olsen, C. A., Expansion of the lysine acylation landscape. Angew Chem Int Ed Engl 2012, 51 (16), 3755–6.

4. Peng, C.; Lu, Z.; Xie, Z.; Cheng, Z.; Chen, Y.; Tan, M.; Luo, H.; Zhang, Y.; He, W.; Yang, K.; Zwaans, B. M.; Tishkoff, D.; Ho, L.; Lombard, D.; He, T. C.; Dai, J.; Verdin, E.; Ye, Y.; Zhao, Y., The first identification of lysine malonylation substrates and its regulatory enzyme. Mol Cell Proteomics 2011, 10 (12), M111 012658.

5. Martin, D. B.; Vagelos, P. R., The mechanism of tricarboxylic acid cycle regulation of fatty acid synthesis. J Biol Chem 1962, 237, 1787–92.

6. Hirschey, M. D.; Zhao, Y., Metabolic Regulation by Lysine Malonylation, Succinylation, and Glutarylation. Mol Cell Proteomics 2015, 14 (9), 2308–15.

7. Du, J.; Zhou, Y.; Su, X.; Yu, J. J.; Khan, S.; Jiang, H.; Kim, J.; Woo, J.; Kim, J. H.; Choi, B. H.; He, B.; Chen, W.; Zhang, S.; Cerione, R. A.; Auwerx, J.; Hao, Q.; Lin, H., Sirt5 is a NAD-dependent protein lysine demalonylase and desuccinylase. Science 2011, 334 (6057), 806–9.

8. Kulkarni, R. A.; Worth, A. J.; Zengeya, T. T.; Shrimp, J. H.; Garlick, J. M.; Roberts, A. M.; Montgomery, D. C.; Sourbier, C.; Gibbs, B. K.; Mesaros, C.; Tsai, Y. C.; Das, S.; Chan, K. C.; Zhou, M.; Andresson, T.; Weissman, A. M.; Linehan, W. M.; Blair, I. A.; Snyder, N. W.; Meier, J. L., Discovering Targets of Non-enzymatic Acylation by Thioester Reactivity Profiling. Cell Chem Biol 2017, 24 (2), 231–242.

9. Nishida, Y.; Rardin, M. J.; Carrico, C.; He, W.; Sahu, A. K.; Gut, P.; Najjar, R.; Fitch, M.; Hellerstein, M.; Gibson, B. W.; Verdin, E., SIRT5 Regulates both Cytosolic and Mitochondrial Protein Malonylation with Glycolysis as a Major Target. Mol Cell 2015, 59 (2), 321–32.

10. Hauser, A.; Penkert, M.; Hackenberger, C. P. R., Chemical Approaches to Investigate Labile Peptide and Protein Phosphorylation. Acc Chem Res 2017, 50 (8), 1883–1893.

11. Herr, R. J., 5-Substituted-1H-tetrazoles as carboxylic acid isosteres: medicinal chemistry and synthetic methods. Bioorg Med Chem 2002, 10 (11), 3379–93.

12. Sureshbabu, V. V.; Venkataramanarao, R.; Naik, S. A.; Chennakrishnareddy, G., Synthesis of tetrazole analogues of amino acids using Fmoc chemistry: isolation of amino free tetrazoles and their incorporation into peptides. Tetrahedron Lett 2007, 48 (39), 7038–7041.

13. Neumann, H.; Peak-Chew, S. Y.; Chin, J. W., Genetically encoding N-epsilon-acetyllysine in recombinant proteins. Nat Chem Biol 2008, 4 (4), 232–234.

