## Supplemental information for "Synthesis and Evaluation of a Stable Isostere of Malonyllysine"

**Table of Contents for Supporting Information**

| | Page |
|---|---|
| Supplementary Figures S1-S2 | S2-3 |
| General materials and methods | S4 |
| Protocol for anti-malonyllysine immunoblotting | S5 |
| Protocol for assessment of amino acid stability | S5 |
| Protocol for anti-malonyllysine dot blot | S6 |
| General synthetic procedures | S7 |
| Synthesis of Fmoc-malonyltetrazole-Lys-OH and Malonyl-NAC | S8-9 |
| $^{1}$H- and $^{13}$C-NMR spectra | S10-15 |
| References | S16 |

**Supplementary Figures**

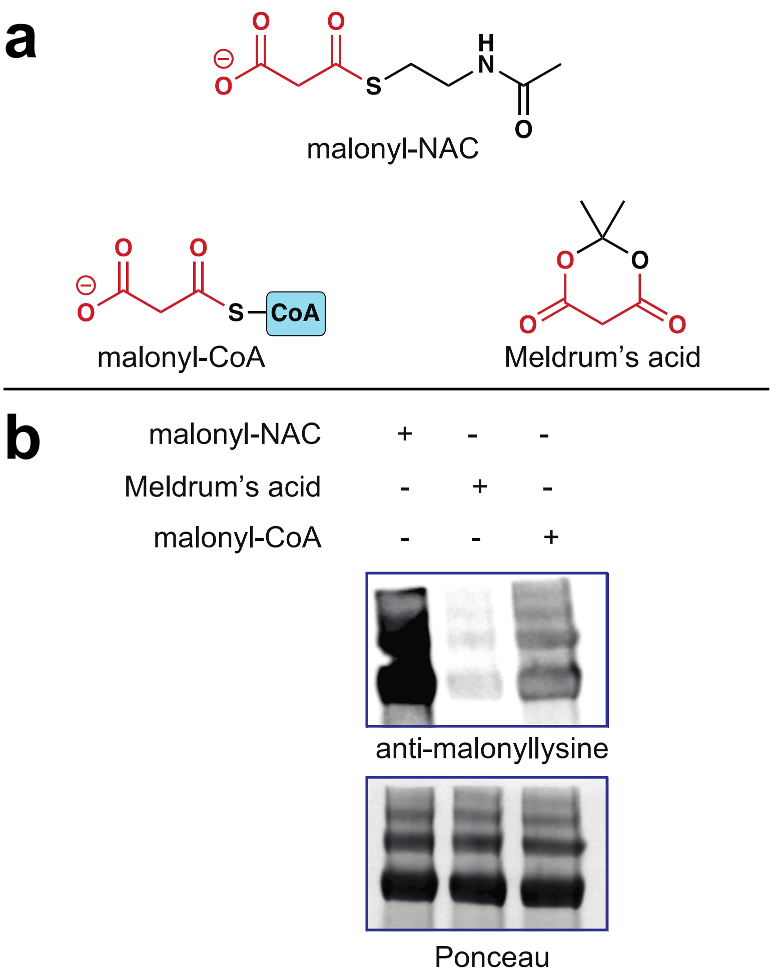


**Figure S1**. (a) Structure of chemical malonylation reagents. (b) Comparison of malonylation of bovine serum albumin (BSA) by chemical malonylation reagents.

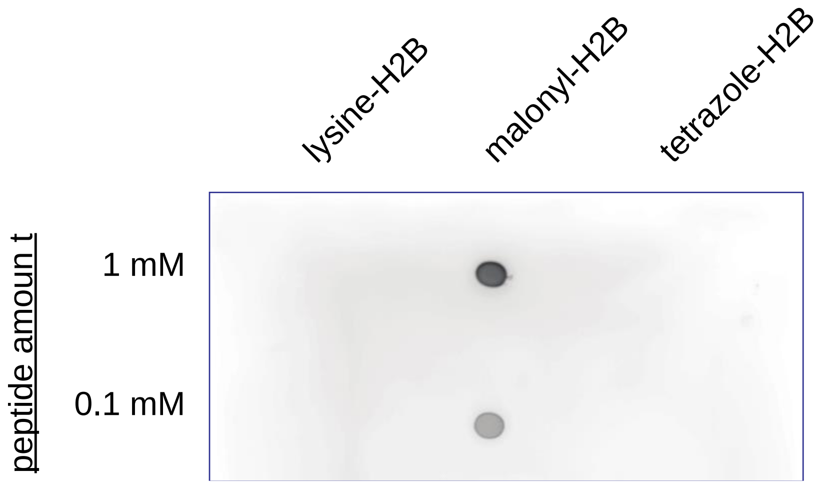


**Figure S2**. A monoclonal anti-malonyllysine antibody (CST #14942) detects malonyllysine peptides more sensitively than tetrazole isostere peptides by dot blot.

### Supplemental Experimental Procedures

#### General Materials and Methods

Non-acetylated bovine serum albumin (#B6917), Meldrum's acid (#210145), and malonyl-CoA (#63410) were purchased from Aldrich. Purity of malonyl-CoA was verified by LC-MS prior to use. Acetyllysine (#9441S) antibodies were purchased from Cell Signaling Technologies (CST) and Cytoskeleton (#AAC01). Anti-malonyllysine antibodies were purchased from Cell Signaling Technologies (CST; #14942S) and PTM Biolabs (PTM-401). SDS-PAGE was performed using Bis-Tris NuPAGE gels (4-12%, Invitrogen #NP0322), and MES running buffer (Life technologies #NP0002) in Xcell SureLock MiniCells (Invitrogen) according to the manufacturer's instructions. SDS-PAGE fluorescence was visualized using an ImageQuant Las4010 Digitial Imaging System (GE Healthcare). Acetyllysine, malonyllysine, malonyltetrazole, and unmodified H2B peptides were purchased from JPT as HPLC-purified peptides (Wang resin) and used without further purification. Analytical LC-MS analysis of malonyllysine and malonyltetrazole decomposition was performed using a Shimadzu LC/MS-2020 Single Quadrupole utilizing a Kinetex 2.6 µm C18 100 Å (2.1 x 50 mm) column obtained from Phenomenex Inc. Runs employed a gradient of 0→90% MeCN/0.1% aqueous formic acid over 4 min at a flow rate of 0.2 mL/min.

**Procedure for Non-Enzymatic Malonylation of Bovine Serum Albumin**[1]

Stock solutions of electrophiles (malonyl-NAC, malonyl-CoA, Meldrum's acid) were dissolved at 20 mM in DMSO. Non-enzymatic malonylation reactions consisted of 40 μg of non-acetylated BSA (10 mg/mL in $H_2O$) and 2 mM electrophile in 50 mM potassium phosphate buffer (pH 8) and 10% DMSO. Reactions were incubated for 6 hours at 37 °C before being quenched by addition of 1x SDS sample buffer. Samples were then heated at 95 °C ("+ heat") or left at room temperature ("- heat") for 5 min as specified, loaded onto 4-12% polyacrylamide gels and analyzed by SDS-PAGE (35 min, 200 V). Following electrophoresis, SDS-PAGE gels were transferred to nitrocellulose membranes for Western blotting using an XCell II Blot Module (30 V, 1 h). Following transfer, blots were blocked for 30 min at room temperature, incubated in primary antibody (anti-malonyllysine, CST #14942, 1:1000 dilution) overnight at 4 °C, washed three times in TBST, and incubated with an HRP-conjugated secondary antibody for 1 h at room temperature (anti-rabbit IgG, HRP-linked CST #7074, 1:1000 dilution). Following an additional three washes, blots were developed via incubation with chemiluminescence detection reagents (Western Blot Detection System, CST) for 1 min prior to image capture using an ImageQuant Las4010 Digital Imaging System (GE Healthcare).

**Analysis of Amino Acid Stability**

For analysis of stability, amino acids (malonyllysine, malonyltetrazole; 100 mM stocks in DMSO) were dissolved in dd$H_2O$ at a final concentration of 1 mM. Samples were incubated for the specified time at the specified temperatures (RT or 90 °C) and analyzed for evidence of decomposition by LC-MS.

**Dot Blot of Malonyllysine and Malonyltetrazole H2B Peptide**

Stock solutions of each peptide were dissolved at 100 mM in DMSO. For dot blot analysis, peptides were diluted to 0.1, 1, 5, and 10 mM in 50 mM potassium phosphate (pH 8). Buffered peptides (1 μL) were then spotted onto nitrocellulose membranes (Novex, Life Technologies # LC2000) and allowed to dry. Membranes were then blocked in StartingBlock (PBS) Blocking Buffer (Thermo Scientific) for 30 min at room temperature, followed by incubation with pan anti-malonyllysine antibodies (PTM Biolabs #PTM-901, 1:2000 dilution; anti-malonyllysine, CST #14942, 1:1000 dilution) for 30 min at 4 °C. Dot blots were washed with TBST three times before addition of HRP-conjugated secondary antibody for an additional 30 min at room temperature (anti-rabbit IgG, HRP-linked [7074], Cell Signaling, 1:1000 dilution). Following an additional three washes with TBST, blots were developed via incubation with chemiluminescence detection reagents (Western Blot Detection System, CST) for 1 min prior to image capture using an ImageQuant Las4010 Digital Imaging System (GE Healthcare).

**General Synthetic Procedures**

Chemicals were purchased from commercial sources (Sigma-Aldrich, Alfa Aesar, and TCI America) and used without further purification unless otherwise noted. Thin-layer chromatography (TLC) was conducted with E. Merck silica gel 60 F254 precoated plates (0.25 mm) and visualized by exposure to UV light (254 nm) or chemical staining. Flash chromatography was performed using normal or reverse phase on a CombiFlash® Rf 200i (Teledyne Isco Inc). $^{1}H$ NMR spectra were recorded at 400 MHz, and are reported relative to deuterated solvent signals. Data for $^{1}H$ NMR spectra are reported as follows: chemical shift (δ ppm), multiplicity, coupling constant (Hz), and integration. $^{13}C$ NMR spectra were recorded at 100 MHz, and are reported in terms of chemical shift (δ ppm). All NMR spectra were standardized to the NMR solvent signal. Analytical LC/MS was performed using a Shimadzu LC/MS-2020 Single Quadrupole utilizing a Kinetex 2.6 μm C18 100 Å (2.1 x 50 mm) column obtained from Phenomenex Inc. Runs employed a gradient of 0→90% MeCN/0.1% aqueous formic acid over 4 min at a flow rate of 0.2 mL/min. High-resolution LC/MS analyses were conducted on a Thermo-Fisher LTQ-Orbitrap-XL hybrid mass spectrometer system with an Ion MAX API electrospray ion source in positive or negative ion mode.

**Procedures for Synthesis of Malonyltetrazole and Malonyl-NAC**

(a) *N*2-(((9*H*-fluoren-9-yl)methoxy)carbonyl)-*N*6-(2-cyanoacetyl)-*L*-lysine or Fmoc-(2-cyanoacetyl)-*L*-lysine-OH

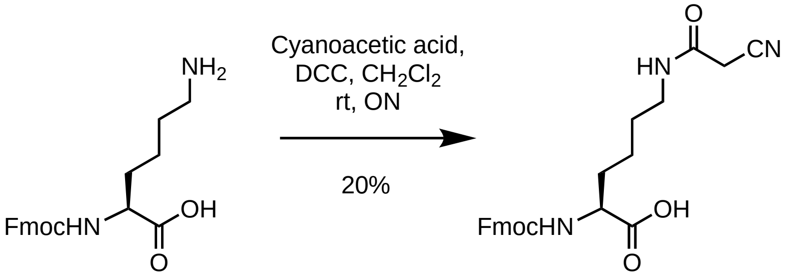


Fmoc-*L*-lysine-OH (3.14 g, 8.55 mmol), 2-cyanoacetic acid (0.87 g, 10.24 mmol), and DCC ( 1.84 g, 8.94 mmol) were dissolved in dry dicholoromethane (20 mL) and allowed to stir overnight at room temperature. The reaction was monitored by Shimadzu LC-MS. After completion of the reaction, reaction mixture was adsorbed on silica gel and purified by flash chromatography (25→ 100% EtOAc:Hexanes) to yield product as amber solid (0.72 g, 20%). $^{1}$H NMR (400 MHz, methanol-$d_4$) δ 7.80 (d, *J*= 7.5 Hz, 2H), 7.67 (t, *J*= 7.9 Hz, 2H), 7.36 (dt, *J*= 32.0, 7.4 Hz, 4H), 4.48-4.29 (m, 2H), 4.22 (t, *J* = 6.8 Hz, 1H), 4.12- 3.96 (m, 1H), 3.36 (s, 2H), 2.91 (t, *J*= 7.4 Hz, 2H), 1.92-1.34 (m, 6H). $^{13}$C NMR (101 MHz, methanol-$d_4$) δ 178.88, 158.07, 145.48, 145.23, 142.62, 128.75, 128.14, 126.23, 126.15, 120.91, 67.72, 57.09, 40.53, 33.47, 28.07, 23.28. MS (ESI+): 436.2, (ESI-): 434.2.

(b) *N*6-(2-(1*H*-tetrazol-5-yl)acetyl)-*N*2-(((9*H*-fluoren-9-yl)methoxy)carbonyl)-*L*-lysine or Fmoc-2-(5-tetrazolyl)acetyl-*L*-lysine-OH2

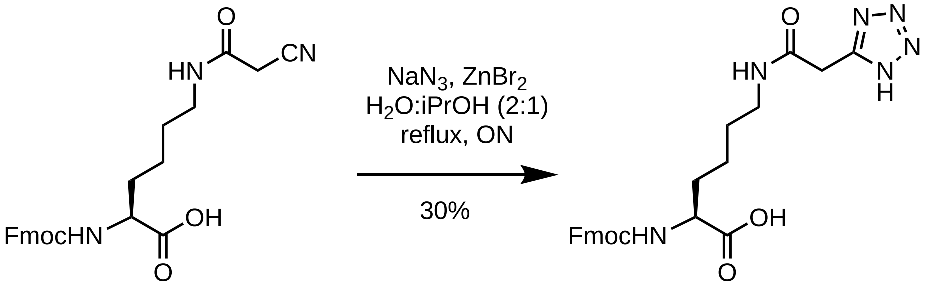


To a stirring solution of Fmoc-(2-cyanoacetyl)-*L*-lysine-OH (0.7 g, 1.02 mmol) in a 2:1 mixture of water and isopropanol (15 mL), sodium azide (0.12 g, 1.82 mmol), and zinc bromide (0.38 g, 1.69 mmol) were added. The reaction was refluxed with vigorous stirring overnight, then allowed to cool to room temperature and quenched with 6M HCl (2 mL). The organic compounds were extracted in ethyl acetate (20 mL). The organic layer was washed with brine (3 x 20 mL), dried over anhydrous $Na_2SO_4$, filtered, and concentrated under reduced pressure. This crude mixture was purified by reverse phase flash chromatography (0→50% $CH_3OH$:$CH_2Cl_2$) to yield the product as dark yellow solid (0.23, 30%). $^{1}$H NMR (400 MHz, methanol-$d_4$) δ 7.80 (d, *J*= 7.4 Hz, 2H), 7.67 (d, *J*= 6.4 Hz, 2H), 7.35 (dt, *J*= 31.9, 7.3 Hz, 4H), 4.36 (q, *J*= 9.8, 9.4 Hz, 2H), 4.30-4.09 (m, 2H), 3.96 (s, 2H), 3.71 (s, 1H), 3.35 (s, 2H), 3.21 (d, *J*= 26.5 Hz, 2H), 1.93-1.30 (m, 6H).$^{13}$C NMR (101 MHz, methanol-$d_4$) δ 175.90, 174.60, 168.78, 158.72, 145.34, 145.17, 142.58, 128.78, 128.16, 126.23, 120.91, 67.91, 55.32, 55.18, 52.69, 48.41, 40.60, 32.29, 32.15, 31.57, 29.69, 24.29, 24.23. MS (ESI+): 479.1, (ESI-): 477.2.

*(c) N*2-(((9*H*-fluoren-9-yl)methoxy)carbonyl)-*N*6-(2-carboxyacetyl)-*L*-lysine or Fmoc-malonyl-*L*-lysine-OH3

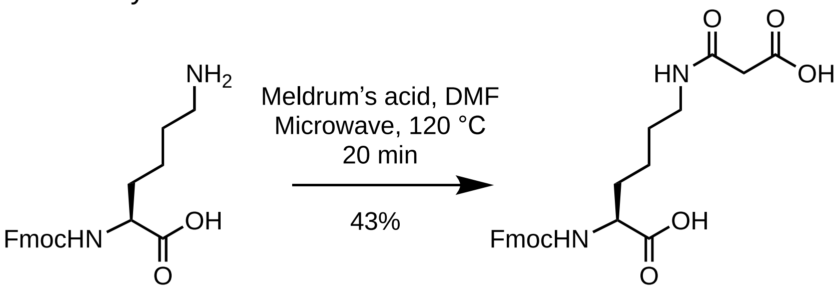


Fmoc-*L*-lysine-OH (0.1 g, 0.25 mmol) and meldrum's acid (28 mg, 0.5 mmol) were dissolved in dry dimethylformamide (2 mL) and subjected to microwave irradiation in a 2-5 mL Biotage microwave vial at 120 °C for 20 min. The reaction mixture was concentrated under reduced pressure and was purified by flash chromatography (0→40% $CH_3OH$:$CH_2Cl_2$) to yield the product as white solid (49 mg, 43%). $^1H$ NMR (400 MHz, $CDCl_3$) δ 7.74 (d, *J* = 7.5 Hz, 2H), 7.58 (q, *J* = 9.5, 8.6 Hz, 2H), 7.42-7.22 (m, 4H), 6.36 (t, *J* = 5.7 Hz, 1H), 5.91 (d, *J* = 8.0 Hz, 1H), 4.47-4.32 (m, 3H), 4.19 (t, *J* = 7.1 Hz, 1H), 3.47 (s, 1H), 3.33-3.10 (m, 2H), 2.96 (s, 1H), 2.89 (d, *J* = 0.7 Hz, 1H), 1.88 (p, *J* = 6.9, 5.5 Hz, 2H), 1.73 (qt, *J* = 18.6, 14.1, 6.3 Hz, 1H), 1.58-1.24 (m, 4H). $^{13}C$ NMR (101 MHz, $CDCl_3$) δ 174.84, 171.69, 163.17, 156.45, 143.95, 143.75, 141.31, 127.80, 127.16, 125.19, 120.04, 67.10, 53.70, 50.57, 47.18, 39.42, 36.83, 32.07, 31.75, 28.69, 22.93, 22.39, 19.84. MS (ESI-): 453.2.

(d) 3-(2-acetamidoethyl)thio-3-oxopropanoic acid or malonyl-NAC free acid3

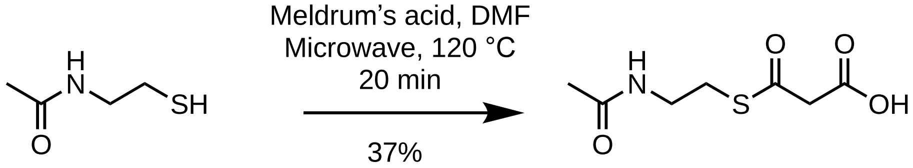


N-acetylcysteamine (0.034 mL, 0.32 mmol) and meldrum's acid (36 mg, 0.64 mmol) were dissolved in dry dimethylformamide (2 mL) and subjected to microwave irradiation in a 2-5 mL Biotage microwave vial at 120 °C for 20 min. The reaction mixture was concentrated under reduced pressure and was purified by flash chromatography (0→40% $CH_3OH$:$CH_2Cl_2$) to yield the product as white solid (24 mg, 37%). $^1H$ NMR (400 MHz, $CDCl_3$) δ 3.66 (s, 2H), 3.50 (q, *J* = 6.1 Hz, 2H), 3.13 (t, *J* = 6.3 Hz, 2H), 1.98 (s, 3H). $^{13}C$ NMR (101 MHz, $CD_3OD$) δ 193.15, 173.51, 169.64, 39.90, 29.65, 22.47. MS (ESI+): 206.1, (ESI-): 204.0.

### $^{1}$H- and $^{13}$C-NMR Spectra

Fmoc-2-(5-tetrazolyl)acetyl-*L*-lysine-OH $^{1}$H NMR

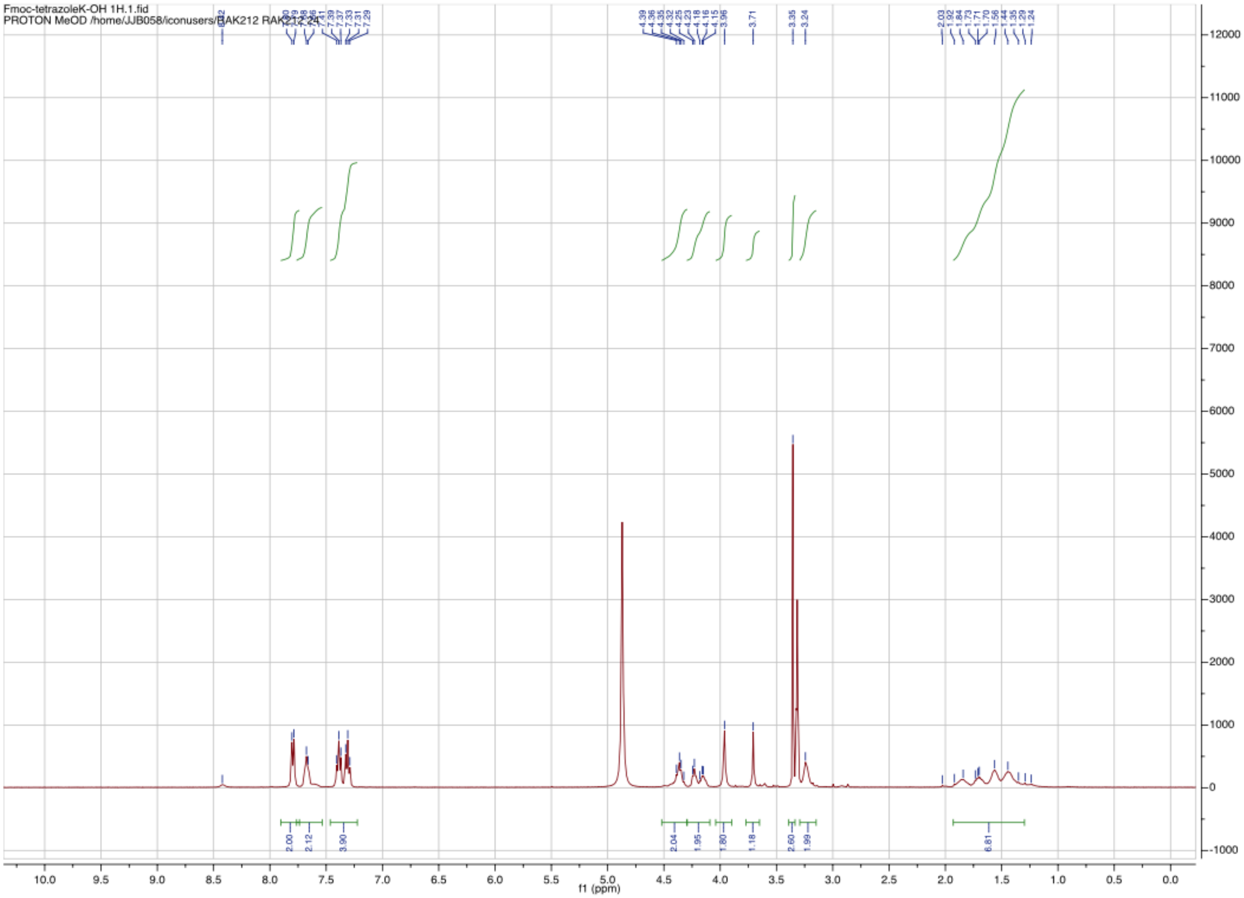

Fmoc-2-(5-tetrazolyl)acetyl-*L*-lysine-OH 13C NMR

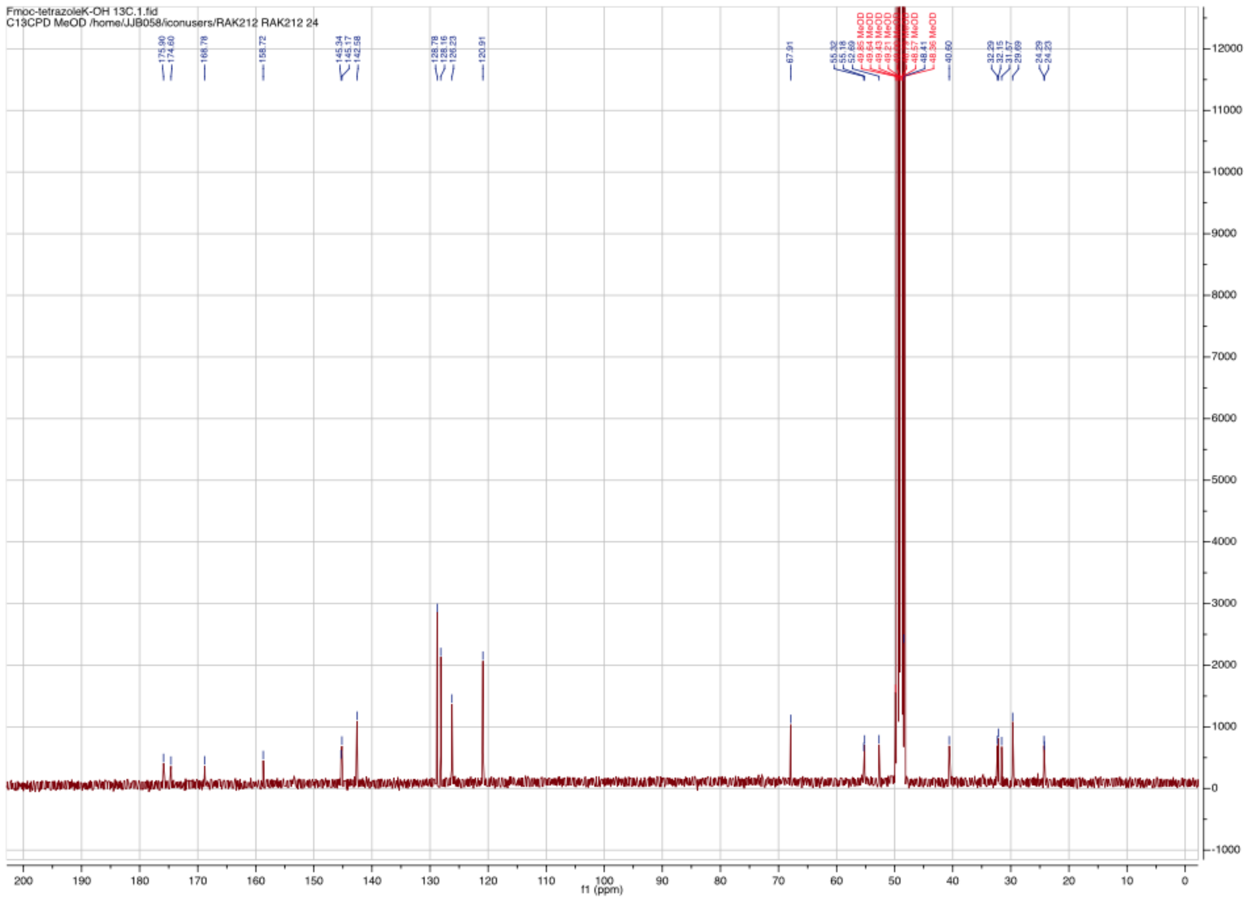

Fmoc-malonyl-*L*-lysine-OH $^1$H NMR

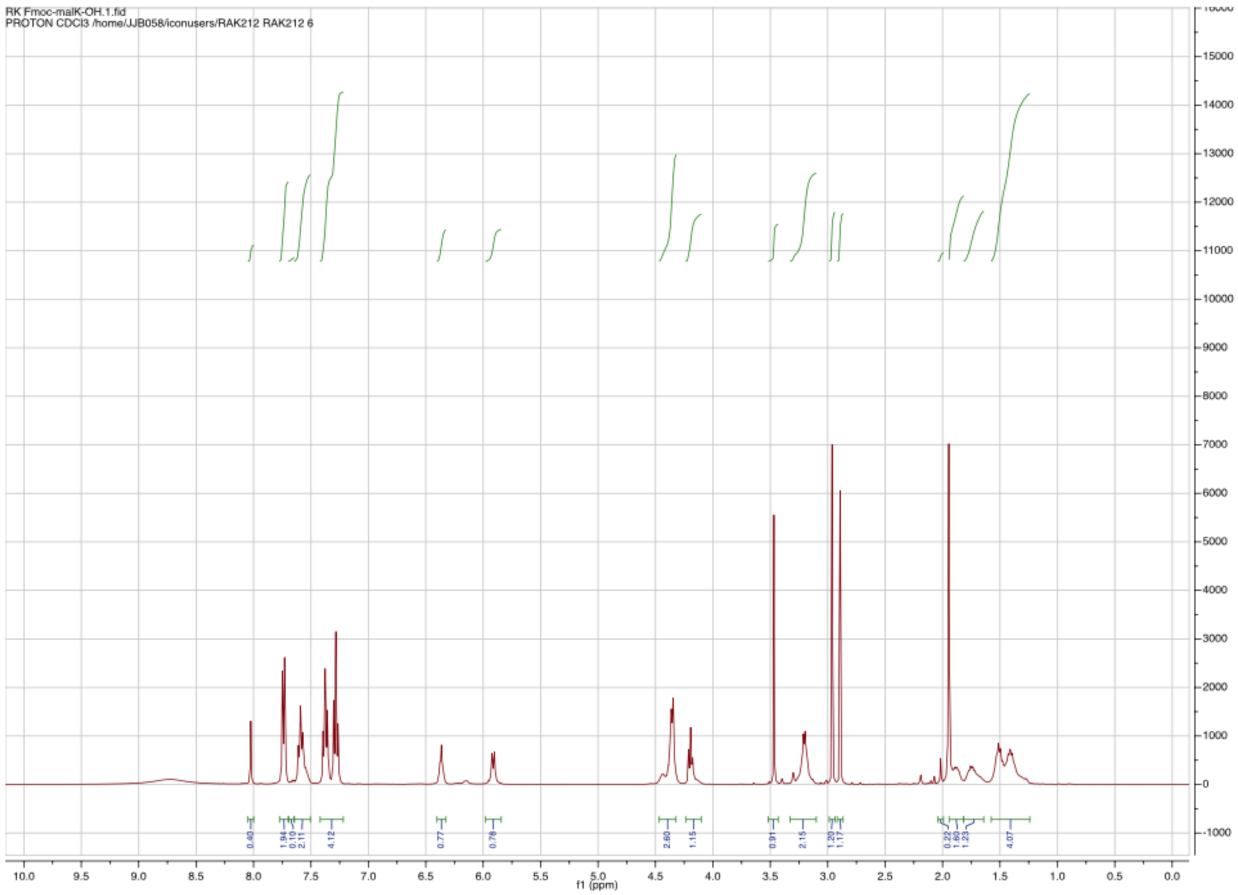

Fmoc-malonyl-*L*-lysine-OH $^{13}$C NMR

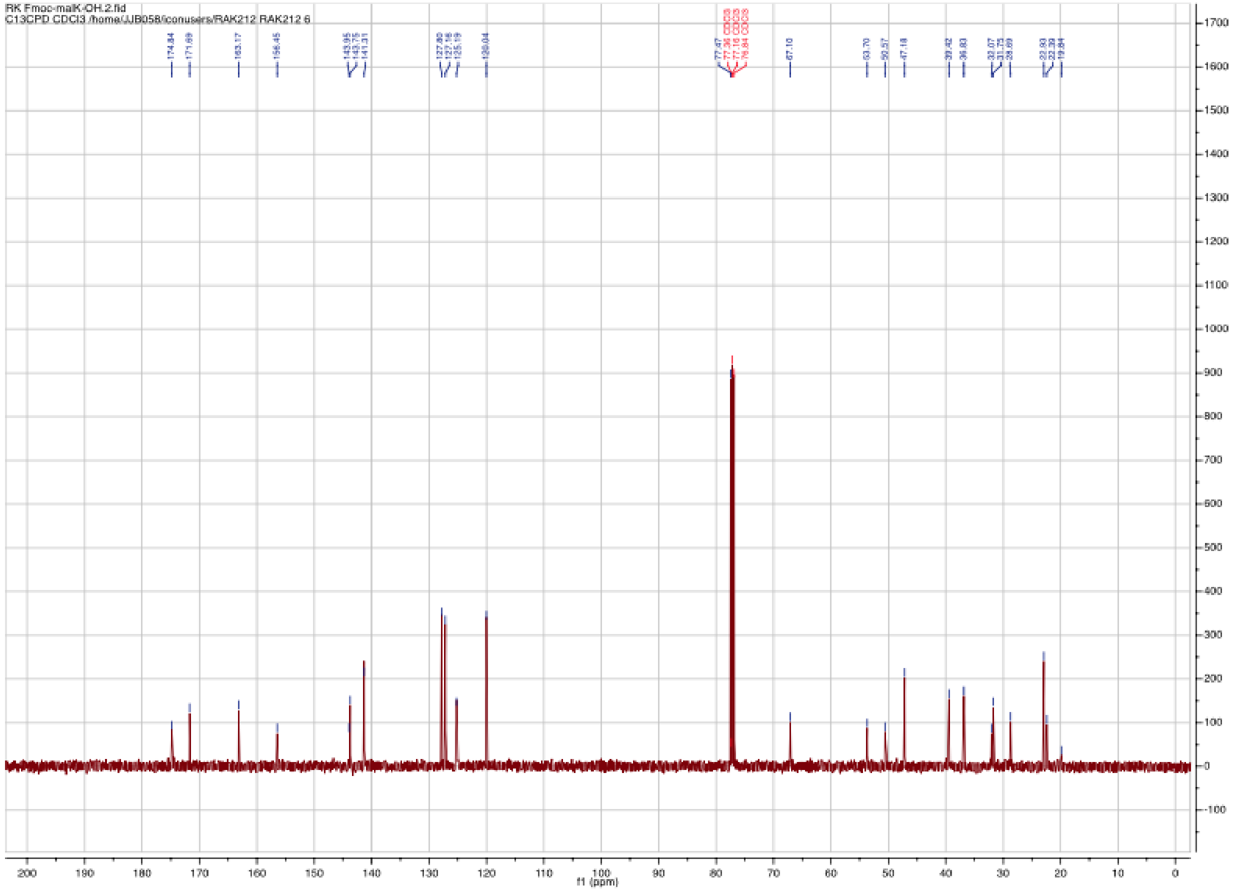

malonyl-NAC free acid $_1$H NMR

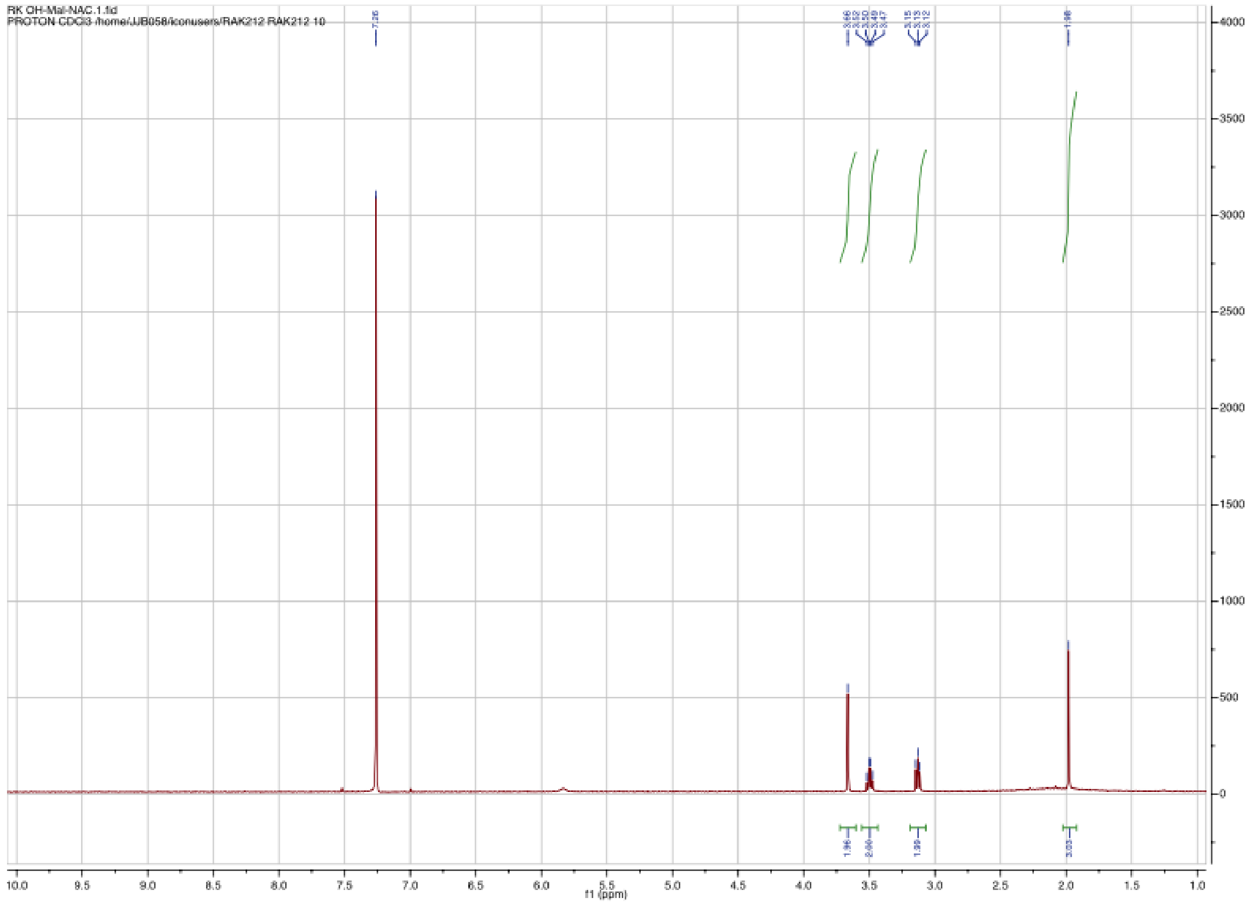

malonyl-NAC free acid 13C NMR

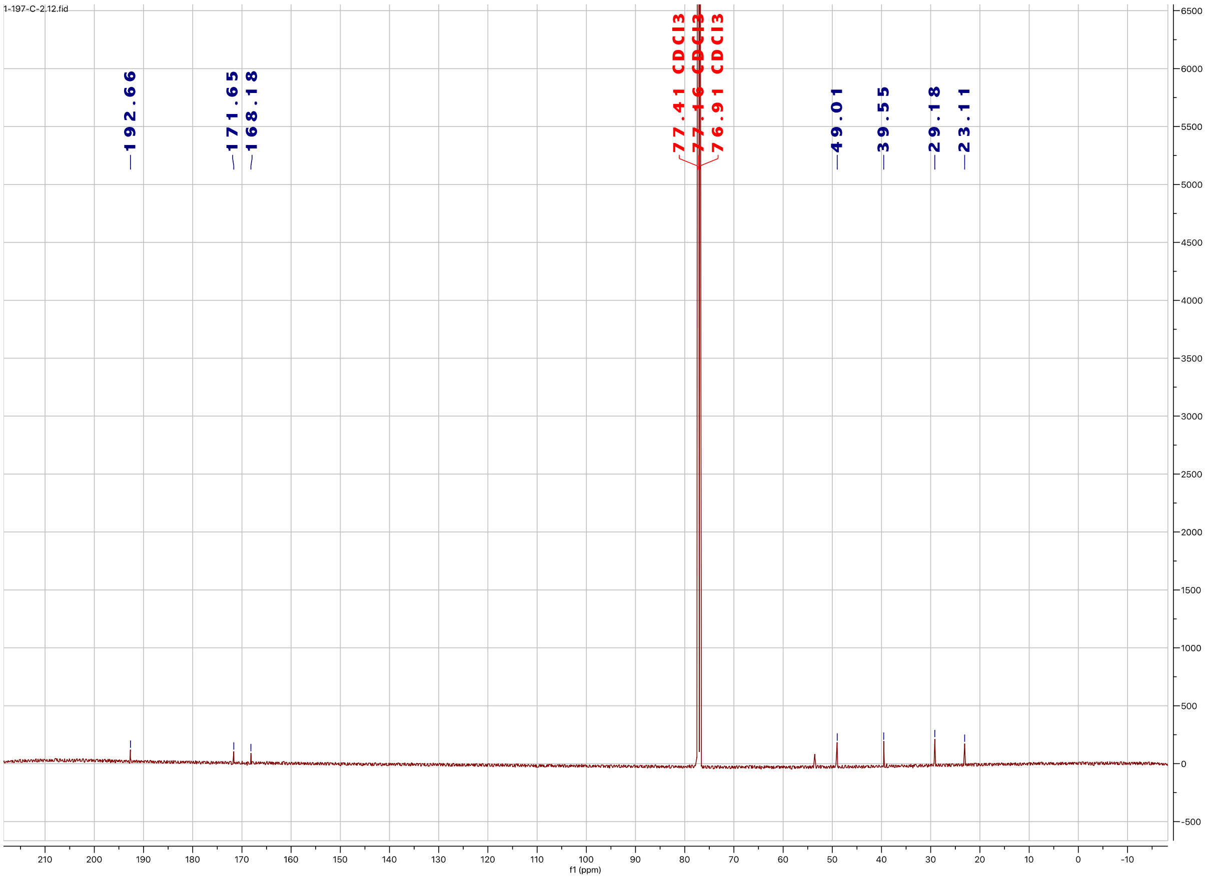

**References:**


1. Kulkarni, R. A.; Worth, A. J.; Zengeya, T. T.; Shrimp, J. H.; Garlick, J. M.; Roberts, A. M.; Montgomery, D. C.; Sourbier, C.; Gibbs, B. K.; Mesaros, C.; Tsai, Y. C.; Das, S.; Chan, K. C.; Zhou, M.; Andresson, T.; Weissman, A. M.; Linehan, W. M.; Blair, I. A.; Snyder, N. W.; Meier, J. L., Discovering Targets of Non-enzymatic Acylation by Thioester Reactivity Profiling. *Cell Chem Biol* **2017,** *24* (2), 231-242.
2. Pallandre, J. R.; Borg, C.; Rognan, D.; Boibessot, T.; Luzet, V.; Yesylevskyy, S.; Ramseyer, C.; Pudlo, M., Novel aminotetrazole derivatives as selective STAT3 non-peptide inhibitors. *Eur J Med Chem* **2015,** *103*, 163-74.
3. Li, Y.; Zhang, W.; Zhang, H.; Tian, W.; Wu, L.; Wang, S.; Zheng, M.; Zhang, J.; Sun, C.; Deng, Z.; Sun, Y.; Qu, X.; Zhou, J., Structural Basis of a Broadly Selective Acyltransferase from the Polyketide Synthase of Splenocin. *Angew Chem Int Ed Engl* **2018,** *57* (20), 5823-5827.
